# A Novel Monocyte-derived Antigen Presenting Cell-T regulatory Cell Axis Contributes to Skin Wound healing and is Impaired in Diabetic Mice

**DOI:** 10.64898/2026.03.04.709590

**Authors:** Jingbo Pang, Brandon Lukas, Rita Roberts, Mark Maienschein-Cline, Yang Dai, Timothy J. Koh

**Author notes:** Address correspondence and reprint requests to Dr. Timothy J. Koh, University of Illinois at Chicago, Center for Wound Healing and Tissue Regeneration, Department of Kinesiology and Nutrition, 1919 W. Taylor Street, Chicago, IL 60612-7246.

## Abstract

Despite a vast literature on the role of macrophages in wound healing, the role of dermal monocyte (Mo)-derived antigen presenting cells (APC) has received scant attention. Using scRNAseq and flow cytometry, we identify a population of APC that is prominent in wounds of non-diabetic mice but is reduced in wounds of diabetic mice. Using adoptive transfer experiments and *Ccr2* knockout mice, we demonstrate that wound APC are derived primarily from Mo and that the diabetic wound environment inhibits differentiation of Mo into APC. We also show that Mo-specific *Irf4* knockout mice exhibit reduced differentiation of Mo into APC, decreased levels of IL27 and numbers of activated Treg cells in wounds. and impaired wound healing. Importantly, adoptive transfer of bone marrow Mo that express *Irf4* into wounds of Mo-specific *Irf4* knockout mice rescued levels of wound APC and activated Treg, as well as wound healing. Local administration of recombinant IL27 into wounds of these mice also rescued levels of activated Treg in wounds, along with wound healing, Together, these findings identify a novel pathway in which IRF4 induces Mo differentiation into APC in wounds, which in turn produce IL27 that activates Treg to promote healing. This pathway is impaired in wounds of diabetic mice, which provides a novel target to improve diabetic wound healing.

## Introduction

Chronic wounds associated with diabetes afflict millions of people and are an increasing problem around the world (1–3). In the United States, individuals with DM face a 25% lifetime risk of developing chronic wounds, with high risk of amputation and a death rate that rivals many cancers. A characteristic of poorly healing diabetic wounds is chronic inflammation, associated with persistent accumulation of inflammatory macrophages (Mφ) (4, 5). Despite a vast literature on the role of Mφ dysregulation in impaired wound healing associated with diabetes, the role of the related family of dermal antigen presenting cells (APC), including classical and monocyte-derived dendritic cells (cDC and MoDC), has received far less attention.

scRNAseq has been used recently to generate comprehensive and unbiased descriptions of myeloid cells and their regulation during normal and impaired skin wound healing (6–10). We identified a wound myeloid cell population that expresses genes associated with antigen processing/presentation, which we identified as APC. These cells express the DC markers *Cd11c*, *Cd40*, *Cd80*, *Cd86* and MHC II genes, but did not express the Langerhans cell marker *Cd207* or the conventional type 1 DC (cDC1) marker *Xcr1* (6). These cells also express genes associated with monocyte-derived DC (MoDC) and conventional type 2 DC (cDC2), including *Cd209a*, *Ahr*, and *Cd301b*, suggesting that the latter DC subsets contribute to wound APC. Previous studies showed that APC can influence wound healing: depletion of epidermal Langerhans cells resulted in either accelerated or delayed wound closure (11, 12), depletion of CD11c+ cells delayed wound closure (13) and MHC II knockout mice exhibit reduced wound collagen deposition and breaking strength (14). However, there is currently little consensus on the role of dermal APC in normal wound healing and a lack of data on diabetic wound healing.

In the present study, we demonstrate that dermal wound APC in are primarily derived from Mo and that the diabetic environment inhibits differentiation of Mo into APC, potentially contributing to impaired healing. We identify the transcription factor IRF4 as a critical regulator of wound APC differentiation, as conditional deletion of *Irf4* in Mo-lineage cells reduces wound APC accumulation and delays wound closure, accompanied by decreased Treg activation and lower IL27 levels in both APC and whole wound homogenates. Local adoptive transfer of *Irf4* expressing CD45.1+ bone marrow monocytes directly into wounds of Mo-specific *Irf4* knockout mice rescues wound APC differentiation, Treg accumulation, and wound healing. In addition, local administration of recombinant IL27 into wounds of these mice also rescues Treg activation and wound healing. Together, our findings define an IRF4-dependent pathway that supports monocyte-derived APC differentiation in wounds, subsequent production of IL27, activation of Treg and wound healing, and further demonstrate that this pathway is impaired in diabetic mice.

## Methods and Materials

### Animals and wound model

C57BL6/J (Non-diabetic, ND), BKS.Cg-*Dock7*^m^ +/+ *Lepr*^db^/J (Diabetic, DB), B6.129S4-*Ccr2*^tm1Ifc^/J (CCR2 knockout), B6.SJL-*Ptprc^a^ Pepc^b^*/BoyJ (CD45.1), B6.129S1-*Irf4*^tm1Rdf^/J (*Irf4*^fl/fl^), and C57BL/6-Tg(Csf1r-cre)1Mnz/J mice were purchased from the Jackson Laboratory (Bar Harbor, ME) and housed for at least two weeks before experiments. *Csf1r*^Cre+^*Irf4*^fl/fl^ mice were generated by crossbreeding B6.129S1-*Irf4*^tm1Rdf^/J with C57BL/6-Tg(Csf1r-cre)1Mnz/J mice. All animal procedures were approved by the Animal care and Use Committee of the University of Illinois at Chicago.

Adult male mice (age 10-12 weeks) were subjected to excisional skin wounding with an 8 mm biopsy punch as described (6). Skin wounds were collected using a 12 mm biopsy punch, then either snap frozen in liquid nitrogen or dissociated by mechanical and enzymatic digestion into single cell suspension as previously described (6).

### Single cell RNA sequencing

Wound cells were pooled from 4 wounds of male ND or DB mice on day 3, 6, and 10 post-injury. Sorted Live CD11b+CD45+Ly6G-cells were processed for scRNAseq using the 10x Chromium Next GEM Single Cell 3’ Reagent Kits (10X Genomics, San Francisco, CA, USA). Libraries were sequenced on HiSeq Sequencing Systems (Illumina, San Diego, CA, USA) with paired-end reads aiming for 100,000 reads/cell.

Ater demultiplexing, cells with >10% mitochondrial gene expression, unique features/gene expression <500 and >10,000, or < 1000 UMI counts were omitted and final 6,109 cells (ND-D3: 961, ND-D6: 1,121, ND-D10: 1,214, DB-D3: 1,334, DB-D6: 894, and DB-D10: 585) were included for downstream analysis (Suppl Fig 1A-1I). After normalization, principal component analysis (PCA) was performed with the 6000 most variably expressed genes (Suppl Fig 1J-1L). Uniform Manifold Approximation and Projection (UMAP) analysis was used for clustering and dimensional reduction. Differentially expressed (DE) gene analysis across all clusters was performed using the Wilcoxon rank sum test to identify marker genes for each cluster with adjusted p-value < 0.05. For cluster composition comparisons, we compared the number of cells in each clustering originating from each sample. Fisher’s Exact test was conducted to construct a 2x2 contingency table based on odds ratio (OR).

**Figure 1.**
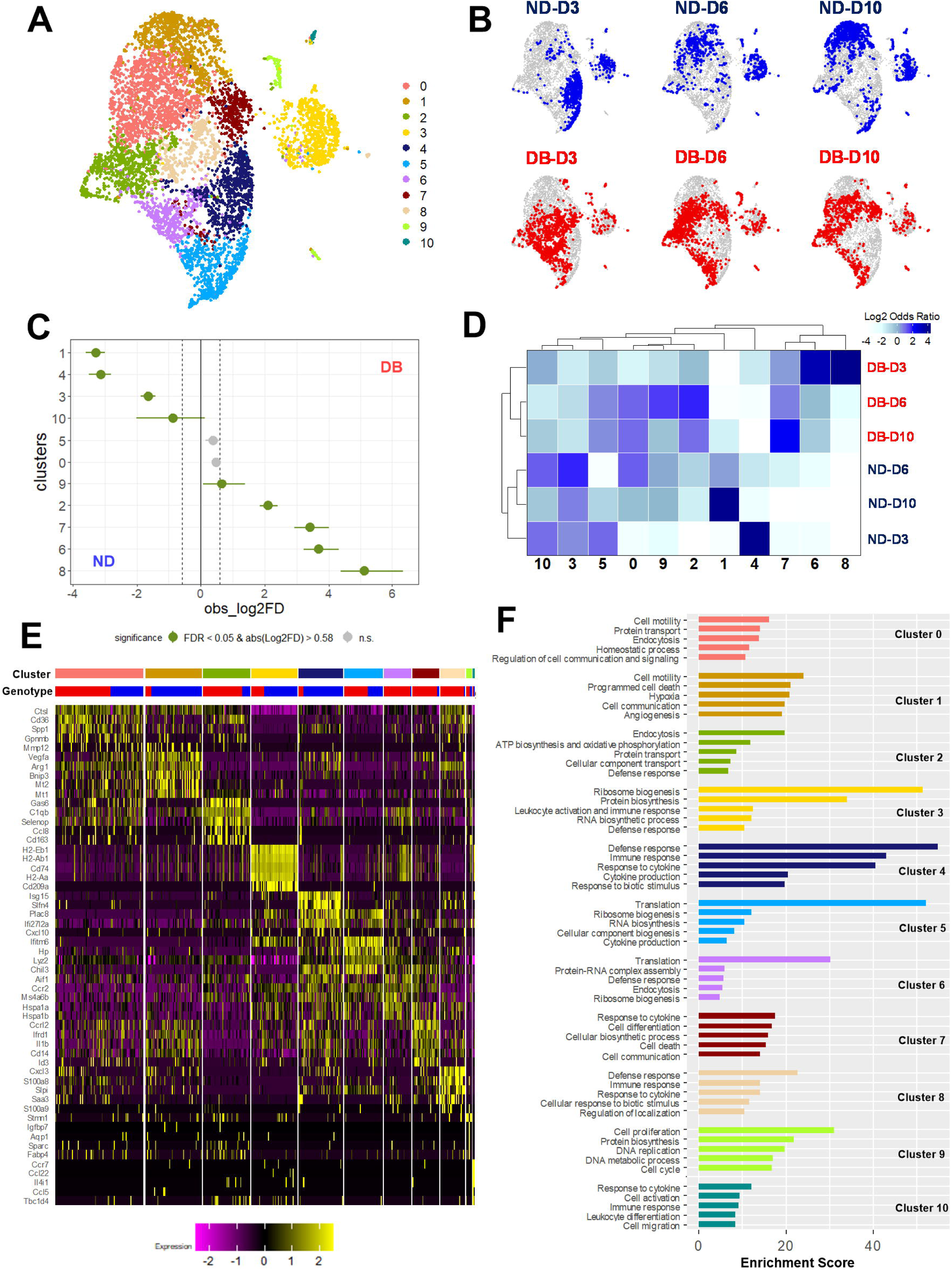
Single cell RNAseq reveals altered non-neutrophilic CD11b+ myeloid cell populations in diabetic versus non-diabetic wounds. Live CD45+CD11b+Ly6G-cells isolated from excisional skin wounds of non-diabetic (ND) and diabetic (DB) mice on days 3, 6, and 10 post-injury and captured using 10x Genomics Chromium platform. (**A**) Top 50 principal components were used for dimensional reduction by UMAP and graph-based clustering analysis identified 11 clusters. (**B**) Feature plots of cell distribution in each cluster indicated a state transition in ND but not in DB wounds. (**C**) Proportion of cells in clusters between the two genotypes was calculated using Single Cell Proportion Test with FDR < 5%. (**D**) Heatmap of cell contribution from ND and DB at each time point to cluster composition based on Log2(Odds Ratio). (**E**) Heatmap displaying the top 5 differentially expressed genes in each cluster. Feature maps showing expression patterns of selected signature genes representing each cluster. (**F**) GO terms which shared similar biological meanings were grouped together in each GO cluster and the top 5 groups were presented and ranked by enrichment score calculated based EASE score of each group member.

Gene Ontology (GO) enrichment analysis of the genes from DE analysis (log2 [fold change (FC)] > 0) from each cluster was performed using The Database for Annotation, Visualization and Integrated Discovery (DAVID) 2021 (FDR < 5%). Functional annotation clustering was conducted using medium classification stringency on all GO terms in each cluster and similar terms were summarized as representative GO terms (Table S2). The overall enrichment score was calculated based EASE score of each group member (15).

### Flow cytometry

Cells were incubated with Zombie NIR™ Fixable Viability Kit (Biolegend, San Diego, CA, USA) followed by Fc blocking by TruStain FcX™ (Biolegend, anti-mouse CD16/32). Next, cells were stained for cell surface markers, and/or intracellular protein using Cytofix/Cytoperm Kit (BD Biosciences). Alternatively, cells were intracellularly labeled with anti-mouse FOXP3 antibody using True-Nuclear™ Transcription Factor Buffer Set (Biolegend) following manufacturer’s instructions. Fluorescence Minus One (FMO) controls were used to calculate the percentage of protein expression within the cell intracellular compartment. All samples were analyzed on Cytek® Aurora cytometer (Cytek Biosciences, Fremont, CA, USA). A complete list of antibodies is presented in Table S1. Data were analyzed using FlowJo (FlowJo LLC, Ashland, OR, USA).

### Adoptive transfer

Bone marrow mononuclear cells from donor CD45.1 mice were enriched by density gradient centrifugation followed by FACS sorting to obtain Zombie-Ly6G-cKit-CD11b+CD115+Ly6C^hi^ Mo on MoFlo Astrios Sorter (Beckman Coulter). Next, 2 million donor Ly6C^hi^ Mos were delivered locally to skin wounds of recipient mice (CD45.2 ND or DB) by intradermal injections on day 2 or 5 post-wounding. Skin wounds of recipient mice were collected 24 hours later and dissociated into single cell suspension for flow cytometry analysis. Adoptively transferred donor Mo-derived APCs were identified as CD45.1+Zombie-CD11b+Ly6G-CD209a+CD74+MHC II+CD64^lo/-^cells.

Alternatively, bone marrow cells from CD45.1 donor mice were enriched by density gradient centrifugation, and monocytes were isolated using the MojoSort™ mouse monocyte isolation kit (Biolegend) following the manufacturer’s protocol. One million CD45.1+ monocytes were locally injected to wounds of *Csf1r*^Cre+^*Irf4*^fl/fl^ mice on day 1, 3, and 5 post-injury. Skin wounds were collected 24 hours after the final injection and processed for flow cytometric analysis (day 6 post-injury).

### Local treatment with IL27 recombinant protein

To assess the function of IL27 in wounds, 360ng recombinant mouse IL27 protein (prepared in 0.1%BSA in 1x DPBS, carrier-free, BioLegend) was injected intradermally around the periphery of each wound of *Csf1r*^Cre+^*Irf4*^fl/fl^ mice at four evenly distributed sites starting immediately after wounding then on day 2 and 4 post-injury (n=4 mice per group). Control mice received injections of an equal volume of sterile 0.1%BSA in 1x DPBS (n=4). On day 6 post-injury, all wounds were harvested, and cells were dissociated from wounds and analyzed by flow cytometry as described above.

### Wound closure and histology

Wound closure measured in digital images of the wound surface taken on day 0, 3 and 6 post-wounding as described (16). Briefly, wound closure was evaluated by measuring wound areas using Fiji Image J and expressing the changes in area at each time point as a percentage of the original wound area ((Day 0 area – Day X area)/Day 0 area)*100 for *Csf1r*^Cre+^*Irf4*^fl/fl^ and *Irf4*^fl/fl^ mice, as well as IL-27-treated *Csf1r*^Cre+^*Irf4*^fl/fl^ mice, and a percentage of the day 1 wound area ((Day 1 area – Day X area)/Day 1 area)*100 for Mo treated wounds of *Csf1r*^Cre+^*Irf4*^fl/fl^ mice, as day 1 was first Mo adoptive transfer in these mice.

Re-epithelialization, granulation tissue area, and collagen deposition measured in hematoxylin and eosin (H&E, Vector Laboratories, Newark, CA, USA) and Trichrome (IMEB Inc., San Marcos, CA, USA) stained cryosections (10μm thick) taken from the center of wounds collected on day 6 post-wounding as described (17). Digital images were taken using a Keyence BZ X710 All in One Fluorescence Microscope with a 2x objective and analyzed using the ImageJ software. The percentage of re-epithelialization was calculated as [(distance traversed by epithelium from wound edges) / (distance between wound edges) ×100]. Three sections per wound were analyzed and data averaged over sections.

### Cytokines

Whole wound tissues from *Csf1r*^Cre+^*Irf4*^fl/fl^ and their control *Irf4*^fl/fl^ mice (day 3 and 6 post-wounding were harvested and homogenized in cold DPBS (10 μL DPBS/mg wound tissue) supplemented with 1% protease inhibitor cocktail (Sigma). Protein levels IL27 and IL10 were determined using a custom BioLegend LEGENDplex™ kit. All samples were analyzed on Cytek® Aurora cytometer (Cytek Biosciences).

### Statistics

Data expressed as mean ± SEM. All data was tested for normality and statistical significance of differences was evaluated by Mann-Whitney test or two-way ANOVA with post-hoc tests. A value of P<0.05 was considered statistically significant.

## Results

### Single cell RNAseq reveals altered myeloid cell populations in diabetic wounds

To investigate dysregulation of non-neutrophilic CD11b+ myeloid cells in DB wounds, we performed scRNAseq on live CD45+CD11b+Ly6G-cells isolated from ND and DB skin wounds on days 3, 6 and 10 post-injury; graph based clustering identified 11 clusters of pooled cells (Fig 1A, Suppl Fig 1). As previously reported (6), wound Mφ exhibited a state transition from pro-inflammatory to pro-healing phenotypes as healing progressed in ND mice (Fig 1B); however, this transition was absent in DB mice.

When analyzing the cell distribution for each cluster, we found that clusters 1, 3, 4, 10 were preferentially populated by ND cells, and clusters 2, 6, 7, 8, 9 by DB cells (Fig 1C). Cluster 4, dominated by cells from early stage ND wounds, differentially expressed genes encoding pro-inflammatory factors, many associated with IFN-signaling pathways (Fig 1D-1F, and Suppl Fig 2). Clusters 6 and 8, mainly composed of cells from early-stage DB wounds, showed high expression of heat shock proteins and S100 calcium binding proteins. Gene ontology (GO) clustering analysis indicated that defense response and cytokine production/response were a feature of all early-stage clusters.

Cluster 1, dominated by cells from late stage ND wounds, showed enhanced expression of antioxidant and pro-healing genes. In contrast, DB cells from day 6 and day 10 wounds were distributed across clusters 0, 2, 5, and 7, and displayed a complex array of phenotypes expressing genes associated with lysosomal function, the complement system, and inflammation. GO analysis indicated several biological processes shared by those clusters, such as cell motility, protein biosynthesis, endocytosis, and immune responses.

Cluster 9 was mainly composed of cells from DB day 6 wounds and showed upregulation of genes associated with cell cycle and cell proliferation, consistent with our previous report of increased Mo/Mφ proliferation in diabetic wounds (18). This cluster also showed enrichment in cell proliferation related pathways in the GO analysis.

Thus, our initial analysis of the scRNAseq data demonstrated a marked cell state transition as healing progressed in ND mice that was absent in DB wounds. Moreover, different inflammatory pathways appear to be activated following skin wounding in cells from DB versus ND mice.

### APC population is decreased in diabetic wounds

Cluster 3 attracted our attention as it was distinct from the other clusters and was over-represented by ND compared with DB cells (Fig 2A). Cluster 3 expressed antigen processing/presentation-associated genes at high levels, particularly *Cd209a*, whereas expression of common Mφ markers was relatively low (Fig 2B and 2C). Thus, we concluded that cluster 3 was a population of wound APC, the accumulation of which is impaired in DB wounds.

**Figure 2.**
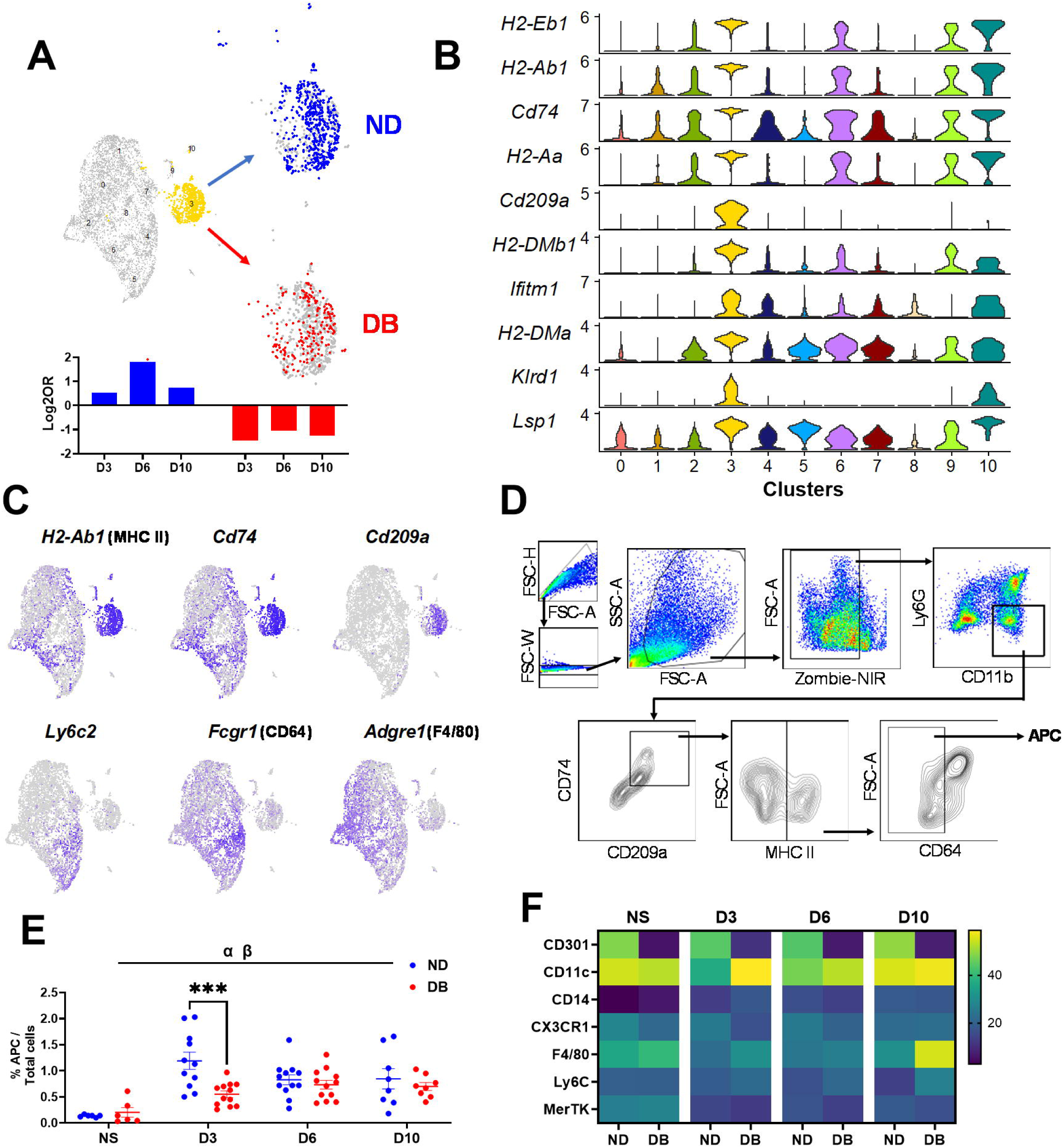
APC significantly decreased in wounds of diabetic mice. (**A**) Feature plots show APC (Cluster 3) over-represented with cells from ND wounds. Log2(Odds Ratio) of cell composition of cluster 3 highlight major contribution from ND day 6 wounds. (**B**) Top 10 signature genes of cluster 3 presented by dot plot to illustrate antigen presentation features. (**C**) Stacked violin plot showing expression patterns of antigen presenting genes and common Mφ markers. (**D**) Gating strategy for identifying APC in skin wounds as Live CD11b+Ly6G-CD209a+CD74+MHC II+CD64^lo/-^. (**E**) Percentages of APC in non-injured skin (NS, ND: n=4; DB: n=6) and wounds on day 3 (n=12/group), 6 (n=12/group), and 10 post-injury (n=8/group) in ND and DB mice from three independent experiments. (**F**) Heatmap of surface protein marker expression in skin wound APC in ND and DB mice. Data are mean ± SEM; ***P < 0.001 between two genotypes by two-way ANOVA. α: significant main effect of time, β: significant main effect of genotype.

To further characterize this APC population, we performed flow cytometry using markers identified by scRNAseq; we defined the APC population as live CD11b+Ly6G-CD74+CD209a+MHC II+CD64^lo/-^cells (Fig 2D). Following injury, both the percentage and number of APC increased in skin wounds in both ND and DB mice. Notably, on day 3, there were significantly lower levels of APC in DB compared with ND wounds (Fig 2E and Suppl Fig 3A). In contrast, pro-inflammatory Mo/Mφ (live CD11b+Ly6G-CD64^hi^Ly6C^hi)^ and mature Mφ (Live CD11b+Ly6G-CD64^hi^Ly6C^int/-^) accumulated at higher levels in DB compared with ND wounds (Suppl Fig 3B and 3C). We also assessed surface expression of several protein markers for Mφ and DC in the APC population. APC expressed Mφ markers CD14, CX3CR1, F4/80, Ly6C, and MerTK at low levels, whereas the DC marker CD11c was expressed at high levels (Fig 2F). CD301, also known as macrophage galactose-type lectin (MGL) and expressed on DC and Mφ (19), was highly expressed in ND APC, with almost no expression on DB APC.

**Figure 3.**
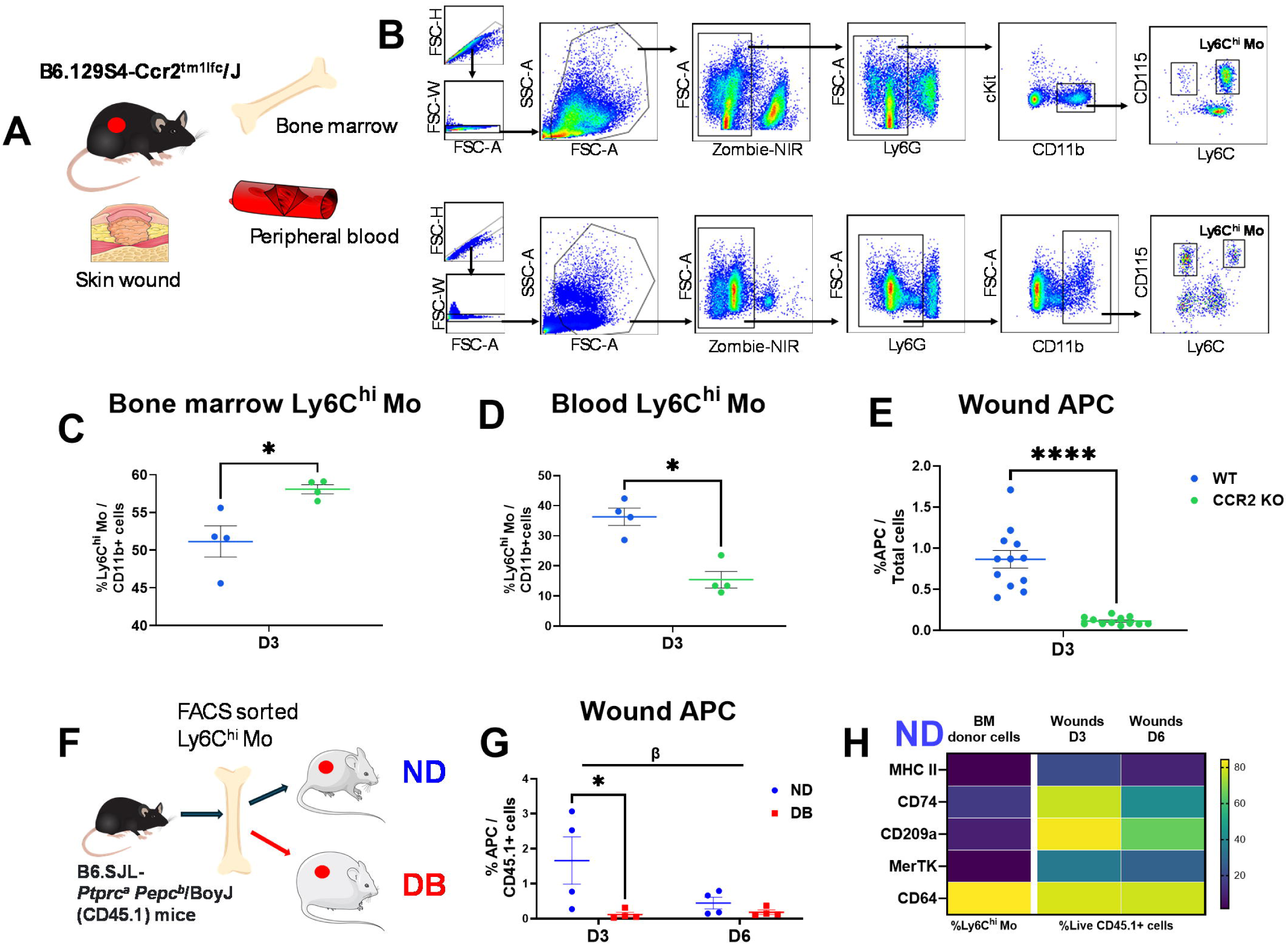
Wound APC originate primarily from infiltrating Mo. (**A**) Schematic of tissue collection from B6.129S4-*Ccr2*^tm1Ifc^/J (CCR2 knockout, KO) mice. (**B**) Gating strategy for identifying Live CD11b+Ly6G-cKit-CD115+Ly6C^hi^ Mo in bone marrow (upper panel) and blood (lower panel). CCR2 deficiency led to increased levels of Ly6C^hi^ Mo in bone marrow (**C**) but decreased levels in peripheral blood (**D**) compared to wild-type (WT) controls (n=4/group from two independent experiments). (**E**) Percentage of APC significantly reduced in CCR2 KO wounds compared to the WT counterparts on day 3 post-injury (n=12/group from three independent experiments). (**F**) Schematic of adoptive transfer of bone marrow Ly6C^hi^ Mo from CD45.1 congenic mice to skin wounds in ND and DB mice. (**G**) Percentages of APC in CD45.1+donor cells 18 hours after transfer to ND vs DB wounds on day 3 and day 6 post-injury. (**H**) Heatmap showing percentages MHC II+, CD74+, CD209a+, MerTK+, and CD64+ cells in bone marrow donor Ly6C^hi^ Mo and CD45.1+ donor cells 24 hours after transfer to ND wounds (BM: n=5; Skin wounds: n=4/group from four independent experiments). Data are mean ± SEM; *P<0.05 and ****P < 0.0001 between two genotypes. β: significant main effect of genotype.

In short, our scRNAseq and flow cytometry data identify a novel population of dermal wound APC, accumulation of which was decreased in DB wounds.

### Diabetic wound environment impairs Mo differentiation into APC

To explore the origin of APC in skin wounds, we first performed experiments with CCR2 KO mice (Fig 3A and 3B). Mo are depleted from peripheral blood in CCR2 KO mice (20), which are often used to study the contribution of circulating Mo to tissue cell populations. As expected, levels Ly6C^hi^ Mo were significantly higher in bone marrow and lower in blood in CCR2 KO mice compared to their wild-type (WT) counterparts (Fig 3C and 3D). In skin wounds collected on day 3, the accumulation of APC was dramatically decreased in CCR2 KO compared to WT mice (Fig 3E), suggesting that infiltrating Mo are a source of APC in wounds.

To determine whether the diabetic wound environment influences Mo differentiation into APC, we performed adoptive transfer of Ly6C^hi^ Mo isolated from bone marrow of CD45.1 congenic mice into wounds of DB and ND mice that express CD45.2 (Fig 3F). Whereas Ly6C^hi^ Mo transferred to day 2 ND wounds differentiated into APC on day 3, this process was significantly impaired in DB wounds (Fig 3G). Very little differentiation occurred when Mo were transferred to day 5 wounds in either ND or DB mice, indicating that differentiation is supported only in the early wound environment. BM Ly6C^hi^ Mo upregulated several APC/Mφ markers after transfer into skin wounds, including MHC II, CD74, CD209a, and MerTK, while expression of the Mφ marker CD64 was maintained at a comparable level or decreased (Fig 3H). A recent study identified DC populations and their precursors within the traditional Mo flow cytometry gate and showed that expression of CD45RB and Flt3 (CD135) could differentiate DC populations from Mo (21). However, CD45RB+CD135+ DC were extremely rare within our BM Ly6C^hi^ Mo population regardless of wounding status in WT mice (Suppl Fig 3D, E), indicating that Mo were indeed the dominant population in our transferred cells.

In short, these data demonstrate that wound APC likely originate from circulating Mo and that the diabetic wound environment significantly impairs differentiation of Mo into APC.

### Transcription factor IRF4 drives wound APC differentiation and improved healing

IRF4 is essential for differentiation of both human and mouse Mo into MoDC (22, 23), and *Irf4* was expressed at relatively high levels in the APC cluster (Fig 4A). To test the role of IRF4 in wound Mo to APC differentiation, we generated mice with IRF4 depleted in CSF1R+ monocyte-lineage cells using *Csf1r*^Cre^*Irf4*^fl/fl^ mice (Fig 4B). Successful Cre-mediated recombination resulted in expression of GFP in IRF4-deficient cells, and we confirmed significantly lower IRF4 protein expression by flow cytometry in GFP+ compared GFP-live CD11b+ myeloid cells in skin wounds (Fig 4C).

**Figure 4.**
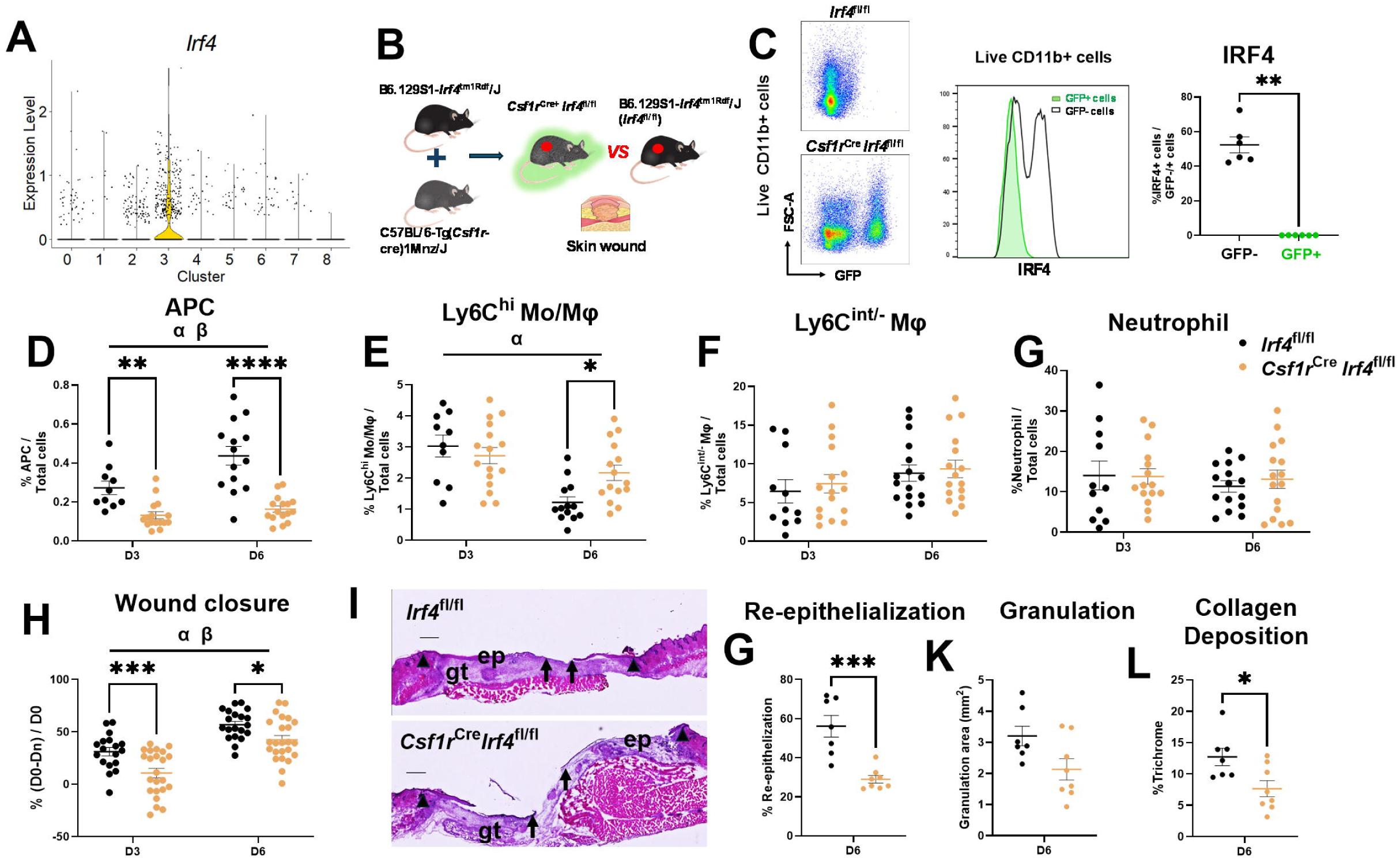
Mo-specific IRF4 knockout reduces wound APCs and impairs healing. (**A**) *Irf4* expression highly enriched in APC cluster compared to other clusters. (**B**) Schematic for crossbreeding B6.129S1-*Irf4*^tm1Rdf^/J (*Irf4*^fl/fl^) with C57BL/6-Tg(*Csf1r*-cre)1Mnz/J mice to generate *Csf1r*^Cre^*Irf4*^fl/fl^ mice. (**C**) Left: Cre-mediated recombination in *Csf1r*^Cre^*Irf4*^fl/fl^ mice confirmed by flow cytometry assessment of Green Fluorescent Protein (GFP) expression. Right: Flow cytometry analysis of IRF4+ cells indicated ablation of *Irf4* in GFP+ cells compared to GFP-cells in skin wounds of *Cfs1r*^Cre^*Irf4*^fl/fl^ mice (n=6/group from two independent experiments). Percentages of APC (**D**), Ly6C^hi^ Mo/Mφ (**E**), Ly6C^int/-^Mφ (**F**), and neutrophils (**G**) in wounds from *Csf1*^Cre^*Irf4*^fl/fl^ and littermate control *Irf4*^fl/fl^ mice on day 3 and 6 post-injury (*Irf4*^fl/fl^: D3, n=11; D6, n=14; *Csf1r*^Cre^*Irf4*^fl/fl^: n=16/group from three independent experiments). (**H**) *Csf1r*^Cre^*Irf4*^fl/fl^ mice showed delayed wound closure compared to *Irf4*^fl/fl^ mice (*Irf4*^fl/fl^: n=18/group; *Csf1r*^Cre^*Irf4*^fl/fl^: n=24/group from three independent experiments). (**I**) Representative images of hematoxylin-eosin–stained (H&E) stained cryosections on day 6 post-injury. ep, epithelium; gt, granulation tissue. Arrows indicate ends of epithelial tongues. Arrowheads indicate the first hair follicle as the original wound, scale bar = 0.5mm. Re-epithelization (**J**) and granulation (**K**) evaluated in H&E stained cryosections of wounds. (**L**) Collagen deposition evaluated by trichrome staining (n=8/group from two independent experiments). Data are mean ± SEM; *P<0.05, **P<0.01, ***P<0.001, and ****P < 0.0001 between two genotypes. β: significant main effect of genotype.

Importantly, APC accumulation was significantly reduced in wounds of *Csf1r*^Cre^*Irf4*^fl/fl^ mice compared to *Irf4*^fl/fl^ controls on both days 3 and 6 post-injury (Fig 4D). In contrast, Ly6C^hi^ Mo/Mφ were increased in *Csf1r*^Cre^*Irf4*^fl/fl^ wounds on day 6 post-injury and Ly6C^int/-^Mφ and neutrophil populations were comparable between the genotypes (Fig 4E-4G), indicating that loss of IRF4 in Mo lead to specific inhibition of differentiation into the APC phenotype.

Associated with reduced APC differentiation, *Csf1r*^Cre^*Irf4*^fl/fl^ mice exhibited significantly delayed wound closure on both days 3 and 6 post-injury compared to *Irf4*^fl/fl^ controls (Fig 4H). *Csf1r*^Cre^*Irf4*^fl/fl^ mice also showed reduced re-epithelialization, granulation, and collagen deposition compared to *Irf4*^fl/fl^ controls in histological analysis of day 6 wounds (Fig 4I-4L). Finally, in the DB wound model, we found lower *Irf4* expression in non-neutrophil myeloid cells and reduced APC accumulation compared to ND counterparts (Suppl Fig 4A and 4B), associated with widely reported impaired healing. Collectively, these data indicate that expression of IRF4 in Mo is required for differentiation to the APC phenotype, and defects in this process leads to impaired healing.

### APC differentiation is critical for the accumulation of Tregs in skin wounds

APC can activate T regulatory cells (Tregs) in response to bacteria and tumor cells (24), and Tregs can promote wound healing (25). Thus, we assessed the impact of reduced APC in wounds of *Csf1r*^Cre^*Irf4*^fl/fl^ mice on Foxp3+ Tregs (Fig 5A). Associated with reduced APC (Fig 4D), wounds in *Csf1r*^Cre^*Irf4*^fl/fl^ mice also contained significantly lower percentages of both Tregs (live CD11b+Ly6G-CD4+Foxp3+) and activated CD25^hi^ Tregs (live CD11b+Ly6G-CD4+Foxp3+CD25^hi^) on day 3 post-injury compared with *Irf4*^fl/fl^ controls (Fig 5B and 5C). Hence, IRF4-mediated APC accumulation in wounds may lead to Treg activation, which in turn, may promote wound healing.

**Figure 5.**
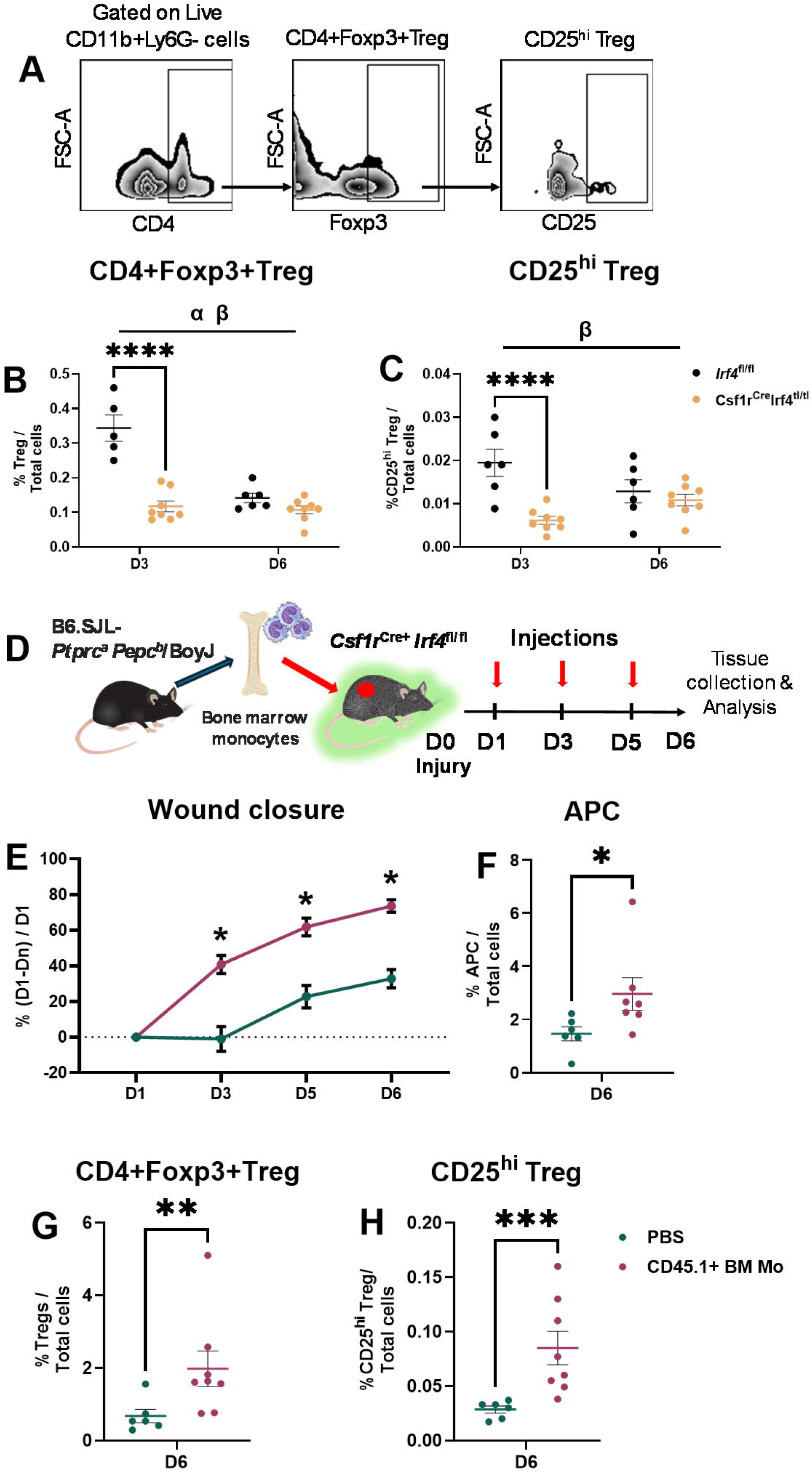
APCs are critical for the accumulation of activated Tregs in skin wounds. (**A**) Gating strategy for identifying Foxp3+Tregs (live CD11b+Ly6G-CD4+Foxp3+) and CD25^hi^ Tregs (live CD11b+Ly6G-CD4+Foxp3+CD25^hi^). Percentages of wound CD4+Foxp3+Tregs (**B**) and CD25^hi^ Tregs (**C**) evaluated on day 3 and 6 post-injury (*Irf4*^fl/fl^: D3, n=5; D6, n=6; *Csf1r*^Cre^*Irf4*^fl/fl^: n=8/group from two independent experiments). (**D**) Schematic of adoptive transfer of bone marrow (BM) Mo from CD45.1 congenic mice to skin wounds in *Csf1r*^Cre^*Irf4*^fl/fl^ mice. (**E**) CD45.1+BM Mo-treated wounds showed accelerated wound closure compared to DPBS-treated control *Csf1r*^Cre^*Irf4*^fl/fl^ mice. Percentages of APCs (**F**), CD4+Foxp3+Tregs (**G**), and activated CD25^hi^ Tregs (**H**) were significantly elevated in wounds in response to CD45.1+BM Mo treatment compared to the DPBS-treatment on day 6 post-injury. CD45.1+BM Mo-treated wounds: n=8/group; DPBS-treated controls: n=6/group from two independent experiments; Data are mean ± SEM; *P<0.05, **P<0.01, ***P<0.001, and ****P < 0.0001 between two genotypes.

To mechanistically assess the impact of APC differentiation on wound Treg activation and healing, we performed an experiment to restore APC levels in wounds of *Csf1r*^Cre^*Irf4*^fl/fl^ mice (Figure 5D). Local delivery of bone marrow Mo from donor CD45.1 congenic mice to wounds of *Csf1r*^Cre^*Irf4*^fl/fl^ mice resulted in significantly accelerated closure rates compared to control mice receiving DPBS (Fig. 5E). This improved healing was accompanied by increased APC accumulation (Fig. 5F) and elevated levels of both total Tregs and activated CD25^hi^ Tregs (Fig. 5G and 5H). Consistent with these findings, DB wounds exhibited significantly reduced accumulation of both Tregs and activated CD25^hi^ Tregs than their ND counterparts (Suppl Fig 4C and 4D), associated with reduced APC and impaired healing. Together, these results confirm the critical role of APC differentiation in regulating Treg accumulation during wound healing.

### IRF4-IL27 signaling axis in APC promotes wound Treg activation and wound healing

Next, we sought to identify a pathway that may mediate communication between APC and Tregs in wounds that leads to improved healing. IL27 has been shown to mediate communication between DC and Tregs in a mouse model of graft versus host disease (26) and IL10 is an important effector of Tregs for wound healing (27). Wound APC from *Csf1r*^Cre^*Irf4*^fl/fl^ mice showed decreased protein levels of IL27 and CD4+CD25^hi^ T cells showed reduced protein levels of IL10 on day 3 post-injury compared to *Irf4*^fl/fl^ controls (Fig 6A and 6B). These reductions were associated with lower IL27 and IL10 protein levels in whole wound homogenates from *Csf1r*^Cre^*Irf4*^fl/fl^ mice on day 6 post-injury (Fig 6C and 6D). Similarly, DB wound APCs expressed lower IL27 protein levels, and Tregs exhibited a trend toward reduced IL10 expression compared to ND controls (Suppl Fig 4E and 4F).

**Figure 6.**
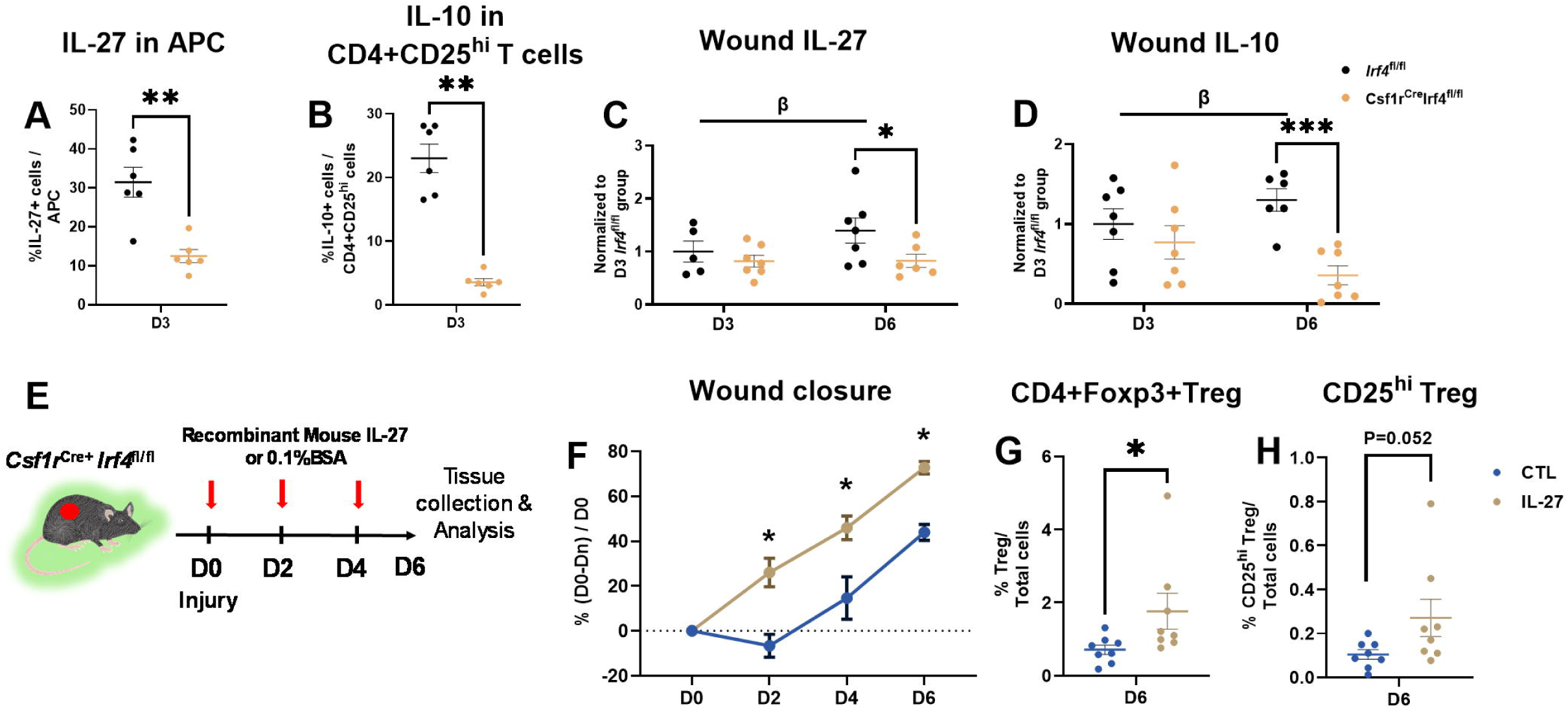
APC-IL27 signaling axis promotes activated Treg accumulation and wound healing. (**A**) Protein expression of IL27 in APC and (**B**) IL10 in CD4+CD25^hi^ T cells (n=6/group from two independent experiments). Due to assay limitations, Foxp3 could not be included in this panel. Wound levels of (**C**) IL27 (*Irf4*^fl/fl^: D3, n=5; n=7 for other groups from two independent experiments and (**D**) IL10 (n=7/group from two independent experiments) measured in wound homogenates on day 3 and 6 post-injury. (**E**) Schematic of local IL27 treatment in skin wounds of *Csf1r*^Cre^*Irf4*^fl/fl^ mice. (**F**) IL27-treated wounds showed accelerated wound closure compared to controls. IL27 treatment increased the percentage of Tregs (**G**), with a trend toward enhanced accumulation of activated CD25^hi^ Tregs in wounds (**H**). N=8 per group; Data are mean ± SEM; *P<0.05, **P<0.01, ***P<0.001, and ****P < 0.0001 between two genotypes; β: significant main effect of genotype.

Finally, we performed a rescue experiment to mechanistically implicate IL-27 in a APC-IRF4-IL27 signaling axis that promotes wound Treg activation and wound healing (Fig 6E). Local administration of recombinant mouse IL-27 protein to wounds of *Csf1r*^Cre^*Irf4*^fl/fl^ mice significantly accelerated wound closure compared to vehicle treated controls (Fig 6F). IL27 administration also increased levels of Tregs (Fig 6G), and induced a trend toward enhanced accumulation of activated CD25^hi^ Tregs in wounds (Fig 6H).

Together, our data indicates that IRF4 is a key promoter of skin wound APC differentiation, which in turn promote healing by activating Tregs via IL-27.

## Discussion

Immune cells orchestrate effective wound healing, but diabetes dysregulates immune cell activity, leading to impaired healing (4, 28, 29). Using scRNAseq, we identified a novel myeloid cell population in murine skin wounds that exhibits features of APC and demonstrated that these cells are derived primarily from Mo using adoptive transfer experiments. Importantly, the diabetic wound environment impaired differentiation of Mo to APC, which may contribute to impaired healing. We also demonstrated that the transcription factor IRF4 served as a key promoter of APC differentiation in wounds; loss of IRF4 in Mo lineage cells disrupted APC differentiation, resulting in impaired production of IL27 and a failure to accumulate activated Tregs, ultimately leading to impaired wound closure. Importantly, these healing defects can be rescued by either the adoptive transfer of wild-type monocytes or local administration of recombinant IL27. In summary, our findings identify a novel pathway in which IRF4-induced Mo-derived APC-activate Tregs via IL27 to promote wound healing.

In this study, we characterized a cluster of wound APC enriched with MHC II genes, *Cd209a* and *Cd74;* this cluster is also prominent in other scRNAseq datasets from different tissues under both steady state and pathological conditions (7, 21, 30). However, these cells have not been characterized in either normal or diabetic skin wounds. These cells also expressed markers of cDC2, MoDC, and Mφ (31, 32), reflecting heterogeneity within the population. Both scRNAseq and flow cytometry analyses demonstrated a significant reduction of APC in DB wounds compared to the ND controls. Our data from CCR2 KO mice and adoptive transfer experiments indicated that APC originate primarily from Mo, and that Mo differentiation into APC is impaired in wounds of diabetic mice, indicating that the diabetic wound environment diabetic environment is hostile to such differentiation.

Previous *in vitro* studies demonstrated that IRF4 is essential for differentiation of both human and mouse Mo into MoDC (22, 23) and *in vivo* studies have highlighted its role in defining DC lineages (33, 34). Our data extend these findings to the wound environment, demonstrating that loss of *Irf4* in CSF1R+ Mo-lineage cells reduced accumulation of APC in skin wounds, whereas accumulation of pro-inflammatory Ly6C^hi^ Mo/Mφ was increased, indicating impaired differentiation of Mo into an APC phenotype, and potentially increased differentiation into an inflammatory Mφ phenotype. These findings indicate that IRF4 is essential for the transition of Mo toward an APC phenotype and its absence leads to preferentially retention of a pro-inflammatory Mo/Mφ fate in skin wound. Additionally, the abnormal shift of immune cell composition also led to defective healing, marked by delayed re-epithelialization and reduced collagen deposition, suggesting that IRF4-dependent APC differentiation is necessary not only for immune regulation but also for normal restoration of injured tissue.

APC promoted Treg activation via production of IL-27 in a mouse model of graft versus host disease (26), and in turn, Treg activation induced release of IL10 to promote the resolution of inflammation and wound healing (27, 35). Indeed, wounds in Mo-specific IRF4 KO mice contained reduced levels of Foxp3+Treg and activated CD25^hi^ Treg on day 3 post-injury, which agrees with a previous report that IRF4 is essential for APC-Treg interaction in skin (36). Importantly, our adoptive transfer experiments mechanistically link IRF4-induced Mo-derived APC, activated Tregs and wound healing, Furthermore, APC that accumulated in day 3 wounds of Mo-specific IRF4 knockout mice expressed less IL27 protein and CD25^hi^ Treg in these wounds expressed less IL10 protein, suggesting impaired activation of local IL27-dependent Treg responses that may contribute to impaired healing in these mice. Our IL27 replacement experiments mechanistically link IL27 to activated Treg accumulation and wound healing.

Interestingly, DB wounds exhibited a similar phenotype as Mo-specific IRF4 KO mice, characterized by reduced APC and Treg accumulation, decreased IL27 and IL10 levels, and impaired healing in these mice. Previous studies have demonstrated that IL27 signaling can promote epithelial cell proliferation and re-epithelialization in skin wounds of non-diabetic mice (37) and can influence Mφ polarization and improve wound healing in the streptozotocin mouse model of diabetes (38). Thus, targeting this pathway may be an appealing approach to improve healing in various contexts.

Our study fills a gap in the understanding of APC heterogeneity and function in skin wound healing, though several limitations remain. First, our scRNAseq data cannot distinguish between resident versus infiltrated cells. Therefore, whereas infiltrating cells likely compose the majority of wound myeloid cells, including APC, we cannot exclude the possibility that resident cells also play a role. Future experiments combining lineage tracing and scRNAseq will be designed to determine the functions of infiltrating versus resident wound APC. Similarly, although our experiments with CCR2 knockout mice and adoptive transfer implicate Mo as precursors of wound APC, we cannot rule out contribution of other potential DC precursors (21). Furthermore, although we focused on cells with an APC phenotype, the functional importance of antigen processing and presentation in wound healing remains unresolved and should be a focus of future study. Finally, the spatial localization of these Mo-derived APCs and their interactions with other wound cells should be more fully elucidated using spatial transcriptomics and downstream experiments.

In summary, our findings advance understanding of the role of Mo-derived cells in wound healing and the dysregulation of these cells in diabetes. We characterized an IRF4-dependent axis in which infiltrating Mo differentiate into wound APC, which produce IL27 that, in turn, promotes Treg activation required for efficient wound healing. Since the diabetic wound environment inhibits this pathway, our findings provide a mechanistic basis for targeting the APC-IRF4-IL27 axis to improve diabetic wound healing.

## Acknowledgments

The authors thank Dr. Giamila Fantuzzi for her input on aspects of presentation of this manuscript. This work was supported by the Northwestern University NUSeq Core Facility.

## Author contributions

JP designed the study, conducted the experiments, performed data and statistical analysis and wrote the manuscript. BL and YD helped design the study, performed scRNAseq data and BITFAM analysis, and helped write the manuscript. RR conducted histology analysis. MMC helped perform scRNAseq data demultiplexing and preparation. TK helped design the study, write the manuscript, and provided all materials.

## Funding

This study was supported by NIGMS through grant R35GM136228 to TJK. MMC was supported in part by NCATS through Grant UL1TR002003.

## Conflicts of interest

All authors have no conflicts of interest to report.

## Notes

### Competing Interest Statement

The authors have declared no competing interest.

