## Supplementary Figure 1-4 for "A Novel Monocyte-derived Antigen Presenting Cell-T regulatory Cell Axis Contributes to Skin Wound healing and is Impaired in Diabetic Mice"

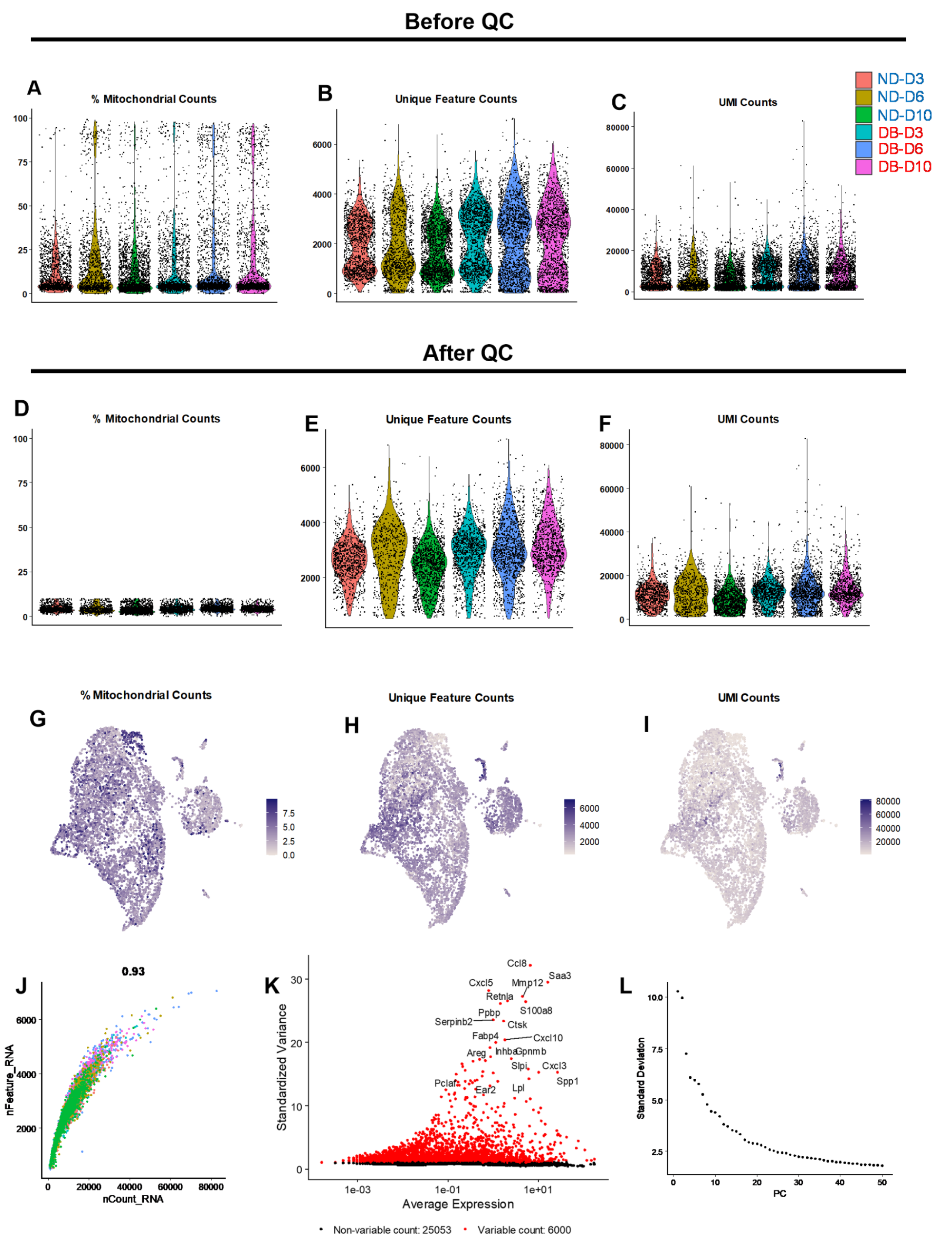


**Supplementary Figure 1. Data processing and quality control**. (A) Quality control filtering (QC): Cells with >10% mitochondrial gene expression, <500 unique features/gene expression, or < 1000 UMI counts were omitted from further analysis. (B) Feature plots showing percentages of mitochondrial, unique features, and UMI count distributions among cells that passed QC . (C) Unique feature counts (nFeature RNA) highly correlated with UMI counts (nCount RNA) post QC. (D) and (E) The top 6000 highly variable genes were used to partition cells along 50 principal components of the dataset.

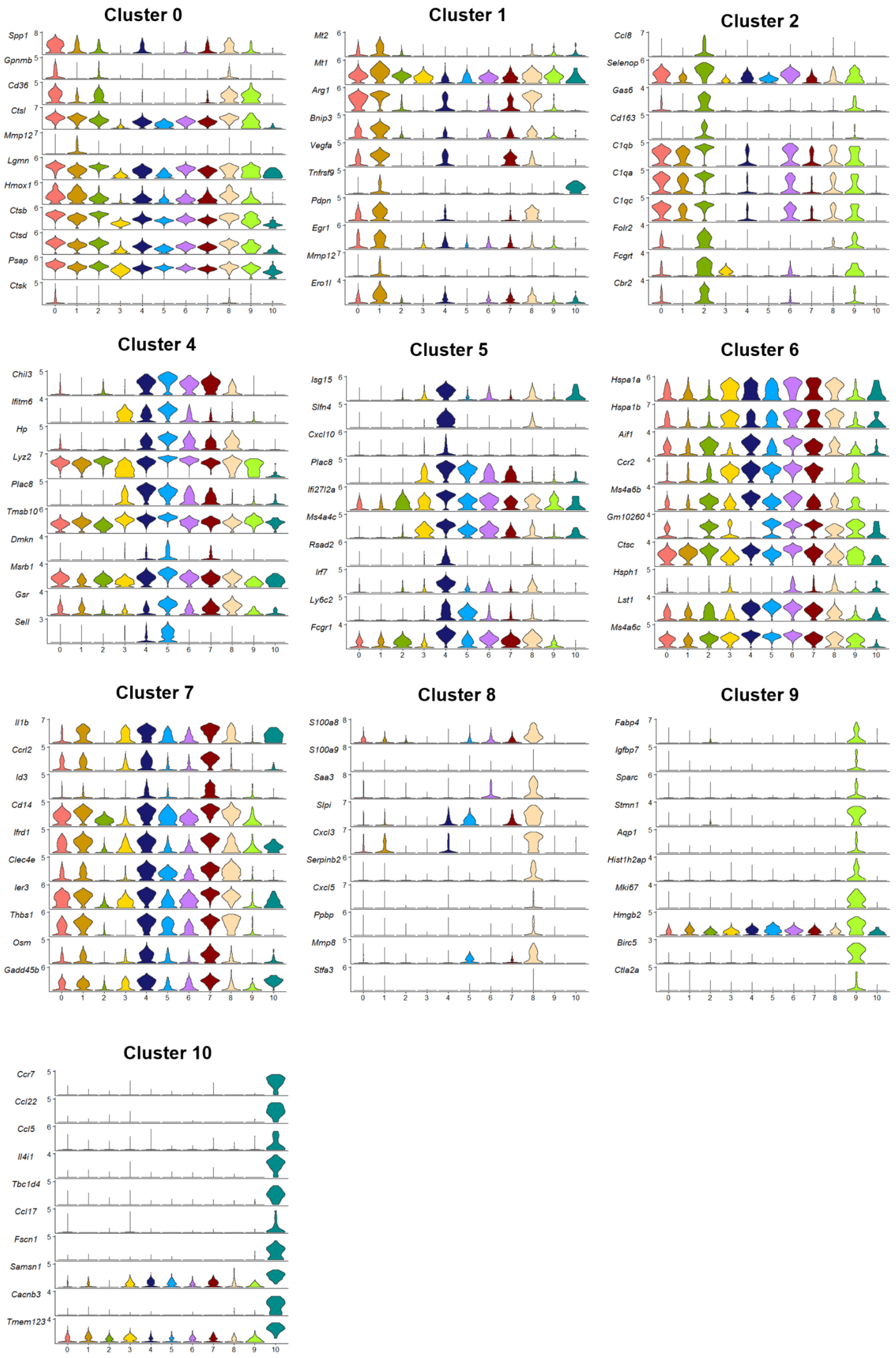


**Supplementary Figure 2. Violin plots of top 10 signature genes of each cluster.** Differential expression across all clusters analyzed using Wilcoxon rank sum test. Violin plots used to illustrate top 10 signature genes for each cluster.

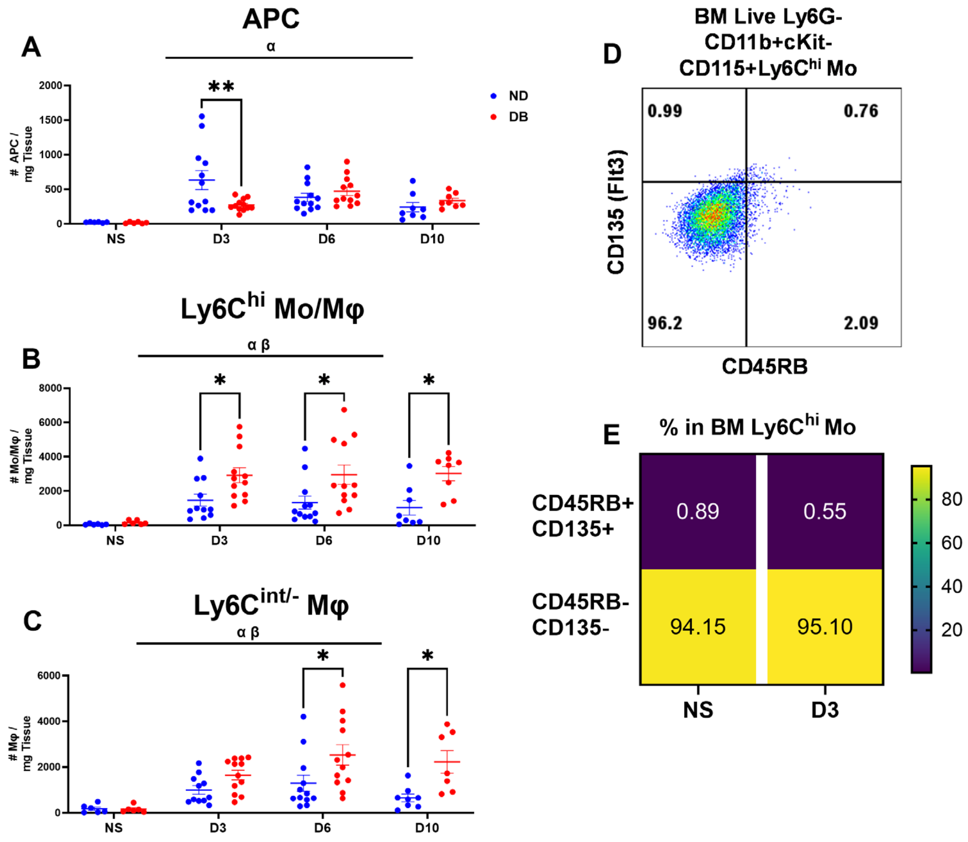


**Supplementary Figure 3. Numbers of inflammatory cells in DB vs ND during skin wound healing.** Numbers of APC (A. Live CD11b+Ly6G-CD209a+CD74+MHC II+CD64$^{lo/-}$), pro-inflammatory Mo/Mφ (B, Live CD11b+Ly6G-CD64$^{hi}$Ly6C$^{hi}$), and mature Mφ (C, Live CD11b+Ly6G-CD64$^{hi}$Ly6C$^{int/-}$) in non-injured skin (NS, n=6/group), and wounds on days 3 (n=12/group), 6 (n=12/group), and 10 (n=8/group) post-injury in ND and DB mice. (D) Representative flow cytometry plot of CD45RB and CD135 (Flt3) expression in bone marrow Live Ly6G-CD11b+cKit-CD115+Ly6C$^{hi}$ monocyte. (E) Heatmap of percentages of CD45RB+CD135+ and CD45RB-CD135- cells within Ly6C$^{hi}$ monocytes in bone marrow collected from non-injured and day3 post-wounding WT mice (n=4/group). Data are mean ± SEM; *$P<0.05$ and **$P < 0.01$ between two genotypes by two-way ANOVA. a: significant main effect of time, b: significant main effect of genotype.

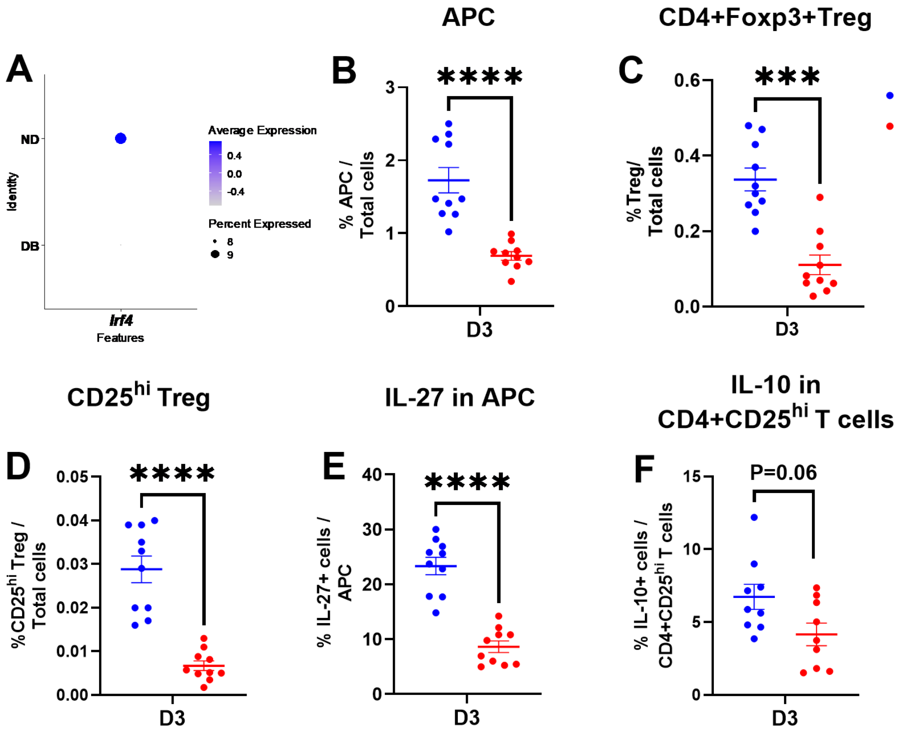


**Supplementary Figure 4. IL27 downregulated in APC and IL10 downregulated in CD4+CD25$^{hi}$ T cells in DB wounds compared to ND wounds.** (A) *Irf4* expression was significantly lower in DB cells compared to their ND counterparts. Percentage of APC (B), CD4+Foxp3+Treg (C), CD25$^{hi}$ Treg (D), IL27 protein expression in APC (E), and IL10 protein expression in CD4+CD25$^{hi}$ T cells (F) in day 3 wounds in ND and DB mice. N=10/group from two independent experiments. Data are mean ± SEM; ****P < 0.0001 between two genotypes by Mann-Whitney test.
